# Continuity between koniocellular layers of dorsal lateral geniculate and inferior pulvinar nuclei in common marmosets

**DOI:** 10.1101/315598

**Authors:** Bing-Xing Huo, Natalie Zeater, Meng Kuan Lin, Yeonsook S. Takahashi, Mitsutoshi Hanada, Jaimi Nagashima, Brian C. Lee, Junichi Hata, Ulrike Grünert, Michael I. Miller, Marcello G.P. Rosa, Hideyuki Okano, Paul R. Martin, Partha P. Mitra

## Abstract

In primates, the koniocellular (K) layers of the dorsal lateral geniculate nucleus (LGN) and the calbindin-rich subdivisions of the inferior pulvinar (IPul) nucleus are considered part of a thalamic matrix system which projects diffusely to superficial cortical layers. Activity in the matrix system is proposed to coordinate oscillatory activity in thalamocortical loops. Further, since both K cells and IPul are involved in visual processing pathways, especially in alternative pathways to visual cortex after V1 lesion in early life (“blindsight”), their functional similarities have been strongly implicated. Here we tested the hypothesis that calbindin-positive K cells and IPul cells constitute a continuous group of cells. By combining immunohistochemistry and a high-throughput neuronal tracing method, we found that both K cells and IPul form reciprocal connections with striate and extrastriate cortices; whereas principal laminae of LGN do not receive inputs from extrastriate cortex and only project sparsely to these areas. Retrograde labelled cells in lateral division of IPul merged seamlessly into the retrograde labelled cells in K layers. These results supported the continuity between LGN K layers and IPul, providing the anatomical basis for functional congruity of this part of dorsal thalamic matrix.

## Introduction

The primate thalamus is a collection of subcortical nuclei that integrate sensory signals and are reciprocally connected to the cortex. The thalamic nuclei are segregated based on histochemical and/or functional differences. Two thalamic nuclei are implicated in visual signal transmission to the cortex, the lateral geniculate nucleus (LGN) and pulvinar (Jones & Hendry, 1989; Saalmann & Kastner, 2011; Sherman & Guillery, 2001). The LGN is seen as a primary relay of retinal signals to cortex (e.g. Preuss, 2007) while the visual parts of the pulvinar are seen as association areas, forming input-output loops with the visual cortices (Shipp, 2003). However, parallel to the view that the LGN and pulvinar have distinct roles, both nuclei are also purported to play a role in synchronizing oscillatory activity of thalamocortical loops and modulate visual cortical activation (Saalmann & Kastner, 2011). It is suggested that both nuclei contain cells that contribute to an underlying matrix of thalamic neurons responsible for synchronized oscillation in cortex (Jones, 2001). Therefore, the question of whether the matrix cells of LGN and visual pulvinar constitute a single population of modulatory thalamic neurons arises.

The proclivity of some thalamic cells to be immunoreactive to calbindin is the main argument for their classification as “matrix” cells, while thalamic cells with parvalbumin immunoreactivity are classed as “core” cells (Jones, 2001). In the primate LGN, three cytoarchitecturally distinct cell classes are identifiable: parvocellular (P), magnocellular (M) and koniocellular (K). P and M cells constitute the principal laminae of the primate LGN, and are immunoreactive for parvalbumin. In contrast, K cells are located between these principal laminae and are immunoreactive for calbindin (Hendry & Yoshioka, 1994; Goodchild & Martin, 1998). K cells receive driving input from widefield retinal ganglion cells and the superior colliculus, and project to superficial layers of V1, V2, the middle temporal area (MT), and the dorsomedial visual area (DM) (Harting *et al.*, 1978; Leventhal *et al.*, 1981; Beck & Kaas, 1998; Hendry & Reid, 2000; Solomon, 2002; Sincich et al. 2004; Szmajda *et al.*, 2008). Furthermore, activity of K cells is synchronized to V1 EEG, thus implicating these cells in the modulatory role of the LGN (Cheong *et al.*, 2011).

The involvement of the pulvinar in visual signal transmission is less well understood than that of the LGN, however its importance in the maintenance of visual perception has been made clear through inactivation studies (Bender & Butter, 1987; Wilke *et al.*, 2010; Purushothaman *et al.*, 2012). Two pulvinar subregions are involved in visual signal modulation: the lateral pulvinar and the inferior pulvinar (IPul). Both the lateral and inferior pulvinar makes reciprocal connections with V1, V2, DM and higher order visual cortices, project to superficial layers of cortex, and contain both calbindin and parvalbumin immunoreactive cells (Benevento & Rezak, 1976; Stichel *et al.*, 1988; Jones & Hendry, 1989; Beck & Kaas, 1998; Gutierrez *et al.*, 2000; Kaas & Lyon, 2007). However, important distinguishing features of inferior pulvinar connectivity are its dense, topographically organised inputs from superior colliculus (Harting *et al.*, 1980; Benevento & Standage, 1983; Stepniewska *et al.*, 2000), evidence for direct input from widefield retinal ganglion cells (Itaya & Van Hoesen, 1983; O’Brien et al. 2001; Warner et al. 2010; Kwan *et al.*, 2018) and projection to MT (Warner *et al.*, 2010).

Due to their similarity in histochemical markers and anatomical organization with respect to their subcortical inputs and cortical connections, we propose that the K cells of the LGN and cells in inferior pulvinar make up a continuous population of thalamic cells primarily involved in regulating the synchronization of corticothalamic loops.

Here we describe anatomical evidence for functional continuity between the LGN K layers and inferior pulvinar by comparing their immunohistochemistry and connectivity with V1 and extrastriate cortices including V2 and DM.

## Materials and Methods

### Immunohistochemistry for calbindin-positive cells

Immunohistochemistry for visualisation of calbindin in the thalamus was conducted on tissue from two male adult common marmosets (*Calithrix jacchus*) acquired from the National Health and Medical Research Council (NHMRC) shared breading facility in Australia. Procedures were approved by the Institutional Animal Experimentation and Ethics Committee at the University of Sydney, and conformed to the Australian NHMRC policies on the use of animals in neuroscience research. Following up to 96 hours in which the marmosets were anaesthetized via intravenous infusion of sufentanil citrate (6–12 μg·kg^−1^·h^−1^, Sufenta Forte, Janssen-Cilag, NSW, Australia) for unrelated physiological experiments, the animal was euthanized (intravenous infusion of 300– 600 mg kg^-1^ sodium pentobarbitone, Lethabarb, Virbac, NSW, Australia) and perfused transcardially with 0.9% saline followed by 4% paraformaldehyde in 0.1 M phosphate buffer (PB, pH 7.4), then by 10% sucrose in PB. The brain was removed and immersed in 20% glycerol for 24–72 hours, then coronally sectioned at 50 μm thickness on a freezing microtome. Sections were initially stained with NeuroTrace blue-fluorescent Nissl stain (1:100; Molecular Probes) for visualisation of nissl substance. Sections were then pre-incubated in PTD for 1 hour at room temperature. The tissue was subsequently incubated with a mixture of primary antibodies (rabbit anti Calbindin and mouse anti Cam Kinase II Abcam) for 2 days under slight agitation. Following this the sections were incubated with secondary antibodies (donkey anti-rabbit Alexa 488 and donkey anti-mouse Alexa 594) for 2 hours. It was not possible to identify any soma labelled for Cam Kinase. However, NeuroTrace blue and Calbindin were evident in all sections.

### Tracing study using the high-throughput neurohistology pipeline

To visualize the thalamocortical connections, we performed tracing study in eight female common marmosets (*Callithrix jacchus*), six of them acquired from the Japanese Central Institute for Experimental Animals and two (M1147, M1148) from Japan National Institute for Basic Biology. Case information is shown in Table 1. All experimental procedures were approved by the Institutional Animal Care and Use Committee at RIKEN and conducted in accordance with the Guidelines for Conducting Animal Experiments at RIKEN Center for Brain Science. For experiments involving participation of Australian researchers, all protocols were approved with a field work license from Monash University and conducted in accordance to the Australian Code of Practice for the Care and Use of Animals for Scientific Purposes.

**Table 1.** Case information and injection sites for marmosets used in tracing study

| Animal ID | Sex | Age | Weight (g) | Tracer | Injection area | Injection site center |  |  |
| --- | --- | --- | --- | --- | --- | --- | --- | --- |
|  |  |  |  |  |  | ML (mm) | AP (mm) | Depth (mm) |
| M820 | F | 6y11mo | 400 | FB | V1 | +1.80 | -8.05 | 0.95 |
|  |  |  |  | AAV-GFP | V1 | +7.25 | -8.15 | 1.10 |
| M822 | F | 7y | 355 | FB | V2 | +6.60 | -6.00 | 0.70 |
| M919 | F | 8y8mo | 345 | FB | V2 | +0.70 | -4.70 | 0.50 |
|  |  |  |  | AAV-GFP | DM | +1.30 | -1.00 | 1.00 |
| M920 | F | 4y7mo | 347 | AAV-GFP | V1 | +0.80 | -7.50 | 0.50 |
|  |  |  |  | AAV-tdTom | DM | +1.00 | -3.20 | 1.00 |
|  |  |  |  | DY | DM | +4.00 | -3.00 | 1.00 |
| M1144 | F | 10y3mo | 353 | FB | V2 | +2.80 | -4.50 | 1.10 |
|  |  |  |  | AAV-GFP | V1 | +3.00 | -8.50 | 1.20 |
|  |  |  |  | AAV-tdTom | V2 | +2.55 | -7.25 | 1.11 |
| M1146 | F | 7y1mo | 394 | FB | V1 | +2.00 | -10.00 | 0.80 |
|  |  |  |  | AAV-GFP | V2 | +8.80 | -7.00 | 2.00 |
| M1147 | F | 4y11mo | 335 | AAV-tdTom | V2 | +3.50 | -5.50 | 0.70 |
| M1148 | F | 4y3mo | 343 | FB | V2 | +5.15 | -4.00 | 1.00 |

Two types of anterograde tracer, AAV-TRE3-tdTomato (AAV-tdTom) and AAV-TRE3-GFP (AAV-GFP), and two types of retrograde tracer, Fast Blue (FB) and Diaminido Yellow (DY), were pressure injected in V1, V2 and DM. Each tracer was delivered using Nanoject II (Drummond, USA) with equal volume at depths of 1200µm, 800µm, and 400µm, controlled with Micro4 (WPI, USA), to fill the entire cortical column. Detailed surgical procedure for tracer injection was described previously (Reser *et al.*, 2009, 2013; Alegro *et al.*, 2017). A summary of all injections is shown in Table 1.

A high-throughput neurohistology pipeline customized for marmosets was adopted to study the mesoscale connectivity (Lin *et al.*, 2018). Briefly, after tracer injection and an incubation period of 4 weeks, the animal was euthanized and perfused. After the brain was fixed with 4% paraformaldehyde, ex-vivo MRI scanning was performed on 6 out of the 8 brains involved in this study. All the brains were then embedded in freezing agent (Neg-50^TM^, Thermo Scientific Richard-Allan Scientific), and cryosectioned coronally at 20 μm using a tape-transfer method (Pinskiy *et al.*, 2013, 2015) customized for marmoset brain. Adjacent sections were treated differently in sequence: either coverslipped directly after dehydration, or processed for Nissl substance, silver staining for myelin, immunohistochemical treatment for cholera-toxin subunit B. Sections processed in the last two ways were not considered in the current study. Therefore 80 μm separated sections were treated the same way. An automatic Nissl staining system (Tissue-Tek Prisma, Sakura, Netherlands) was used to stain those sections processed for Nissl substance with Cresyl Violet. With the design of the custom tape-transfer method, different types of sections were processed in batches. The process of mounting coverslips was also automated using Tissue-Tek Glas, GLAS-g2-S0 (Sakura, Netherlands).

All brain sections were scanned in a high-throughput fashion in Nanozoomer 2.0 HT (Hamamatsu, Japan) as 12-bit RGB images, with a resolution of 0.46 μm/pixel. Unstained sections were scanned for fluorescence using FITC/TX-RED/DAPI as the excitation light filter; while the Nissl-stained sections were scanned with bright-field imaging. The RAW images were processed in a high-performance computational infrastructure (Lin *et al.*, 2018). Images of individual brain sections were isolated and compressed into JPEG2000 format for economic data storage and subsequent analyses.

### Tracer-labelled neuron detection

An automatic cell detection routine was developed to locate retrograde tracer-labelled cells in fluorescent sections. In short, a mask of the brain section was calculated to remove the glass slide background. A Mexican hat filter was applied to quench background and foreground noises. Color and intensity thresholds were applied to weakly filter the pixels. A series of morphological operations were performed to identify individual cells and clusters of cells. Clusters of cells were cut using an fast, unsupervised method (Pahariya *et al.*, 2018). Coordinates of the cell centroids were recorded to represent individual cell location in the brain section. The results were uploaded to the web portal. Manual proofreading was performed to correct false detections.

Anterograde tracer-labelled axon terminals were identified manually for each brain section as areas with fluorescent label intensities higher than an empirically defined threshold.

### Cross-modality registration

In order to identify the brain region where the tracer-labelled cell body or axon terminals were detected, it was necessary to overlay the fluorescent section with its adjacent Nissl section, where the brain region segmentation was based on. A computational routine was established to perform this process automatically for all brain sections (Lee, Tward, *et al.*, 2018). Briefly, in 6 animals where *ex vivo* pre-sectioning MRI of the brains were available, the Nissl stack was reconstructed by a series of jointly optimized rigid motions using the MRI as a guide, and the resulting reconstructed stack was diffeomorphically registered to the marmoset brain atlas (Hashikawa *et al.*, 2015). In the 2 animals (M820, M822) where MRI guidance was unavailable, reconstruction was performed using the atlas as a shape prior and accounting for intrinsic shape differences (Lee, Lin, *et al.*, 2018). The fluorescent sections were then aligned and registered to the reconstructed Nissl stack using a similar joint method, also over a series of rigid motions. The calculated transformation was applied to the cell coordinates and terminal areas, so that the tracer-labelled neurons were mapped onto the segmented brain regions on Nissl sections (Fig. 2C,F,I, Fig. 4A).

**Figure 1.**
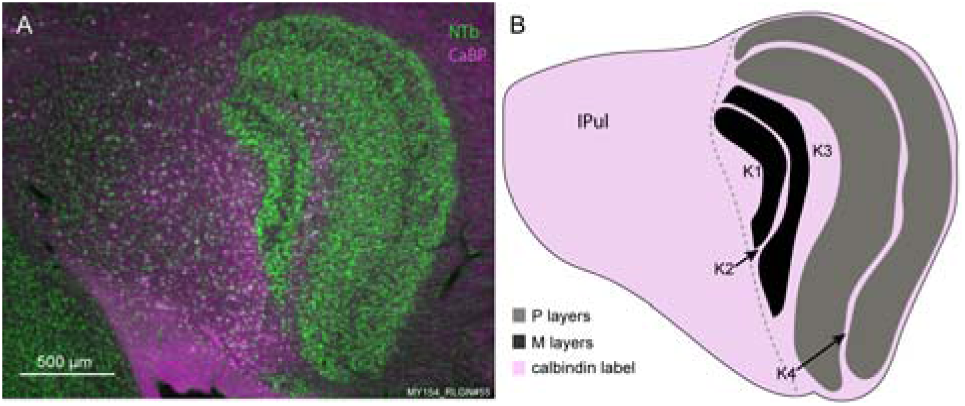
**A.** Photomicrograph of a coronal section through LGN and pulvinar. Soma stained for Nissl substance using NeuroTrace blue (NTb) are shown in green. Soma stained for Calbindin (CaBP) are shown in magenta. **B.** Drawing outlining the LGN laminae and IPul in A. The region with calbindin labelling is shown in light magenta. The grey dash line shows the proposed border between LGN and IPul. IPul: inferior pulvinar; P layers: parvocellular layers; M layers: magnocellular layers; K1-4: koniocellular layers 1–4.

**Figure 2.**
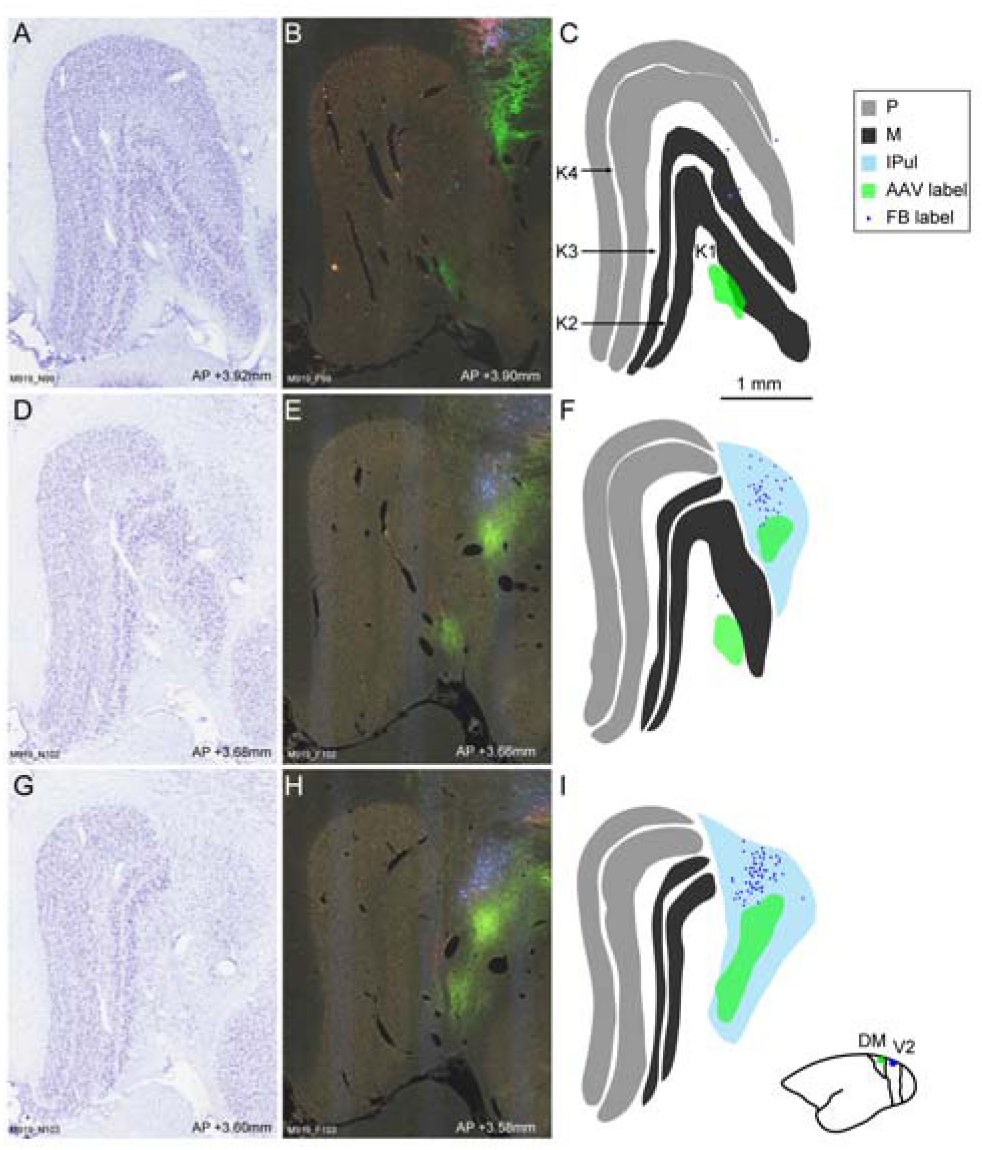
Series of sections through the LGN and IPul of case M919 showing retrograde label and anterograde terminals. **A**. Nissl stained section showing LGN. **B**. Fluorescent section adjacent to A, showing retrograde Fast Blue labeled cell bodies and anterograde AAV-GFP labeled axon terminals. **C**. Annotated brain regions based on A, with locations of FB-labeled cell bodies (blue dots) and AAV-GFP-labeled axon terminals (green area) from B. **D-F**. Adjacent Nissl-stain, fluorescent, and brain region annotation images for a more caudal section, where IPul was visible. **G-** A further caudal set of sections. **Inset**: An illustration of the injection positions on the brain atlas injection locator. Blue dot: Fast Blue injection site. Green dot: AAV-GFP injection site. Abbreviations the same as in Fig. 1.

### Connectivity quantifications

For both anterograde and retrograde tracing cases, the existence of each connection for every individual case was indicated as a dummy variable. The consistency of existence of each connection was obtained by averaging the dummy variables across all related cases and reported in percentage (Fig. 3A, B).

**Figure 3.**
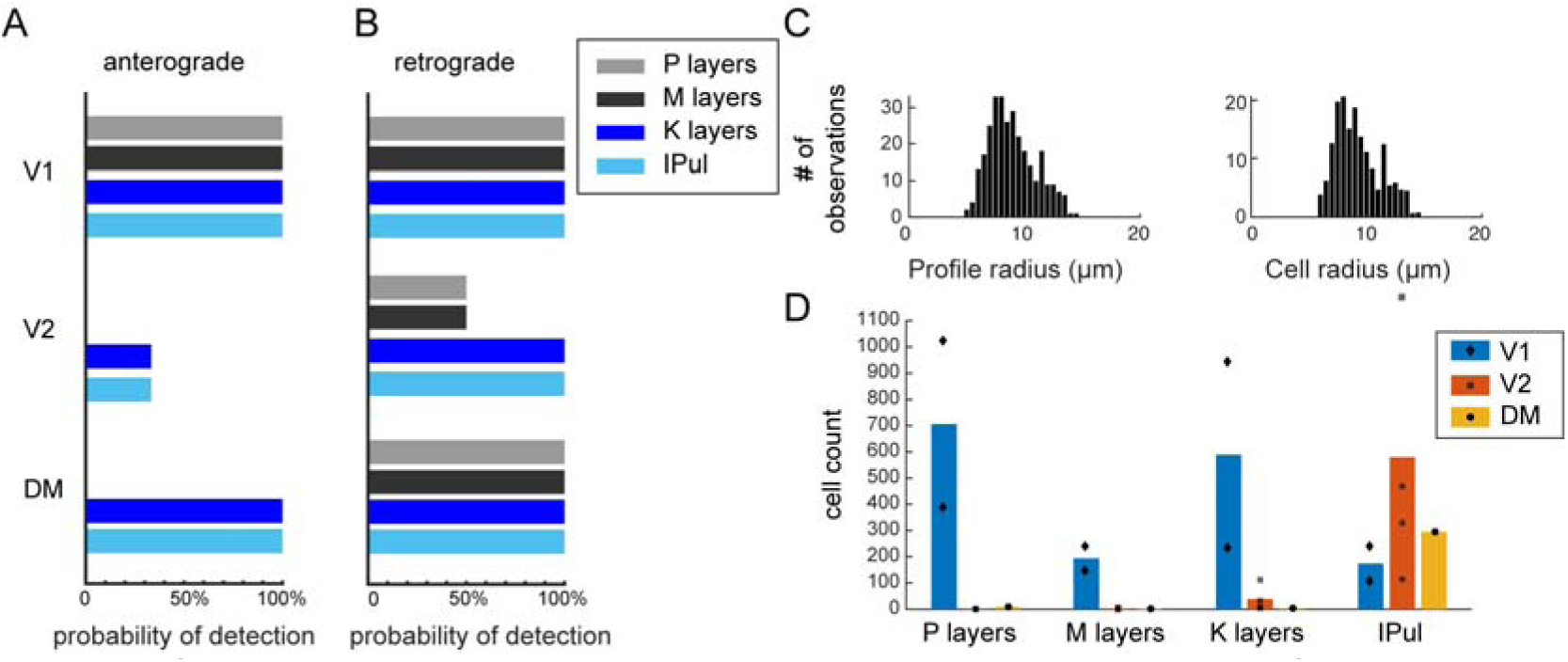
**A**. Probabilities of corticothalamic projections from visual cortices to LGN layers or IPul based on anterograde tracing results across all cases. A probability of 100% means the cortical projection to this area exists in all cases. Injection cases: V1: N=3; V2: N=3; DM: N=2. **B**. Probabilities of thalamocortical projections from LGN layers or IPul to visual cortices based on retrograde tracing results across all cases. Injection cases: V1: N=2; V2: N=4; DM: N=1. **C**. The histogram of cell profile area from cell detection (left) and the histogram of cell area (right) estimated from profile distribution. **D**. Average number of cells detected in LGN layers and IPul across retrograde tracing cases. Cell count for each case was plotted as diamonds for V1 injection, squares for V2 injection, and circles for DM injection. In one case of V2 injection, the corrected cell count mounted up to 1408, and is represented by the square marker above the labelled range of y-axis.

For retrograde tracing cases, fluorescent labelled cells were counted within each brain region. Profiles of cell body were detected using the method described above (“Tracer-labeled neuron detection”). True number of cells in each section was estimated based on the distribution of profile area and the section thickness using a recursive reconstruction method (Rose and Rohrlich 1988). From 298 profiles sampled from two 20-μm sections with FB labelled cells, the profile-to-cell ratio was estimated to be 1.75 (Fig. 3C). For the gap of 80 μm between sections, we assumed linear relationship of labelled cell numbers across sections. Therefore the true cell number within each brain region was estimated as

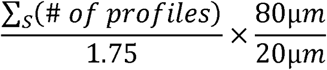

where *S* is the total number of brain sections containing a particular brain region. In Fig 3D, we reported both average number of cells across all cases (filled bars), and the cell count for individual cases (dark grey markers).

The rostral-caudal transition of FB labelled cell bodies between LGN and pulvinar was shown as the accumulated cell numbers for each individual coronal section. To align fluorescent stacks for all cases, we set the most rostral section with IPul as the baseline section d_ip_=0. Then for all cases with the same injection region (V1 or V2), the coronal sections with the same relative distance to this baseline section were surveyed. The cell number at each distance was the accumulated number of cells within each brain region for all the surveyed sections (Fig. 4B).

**Figure 4.**
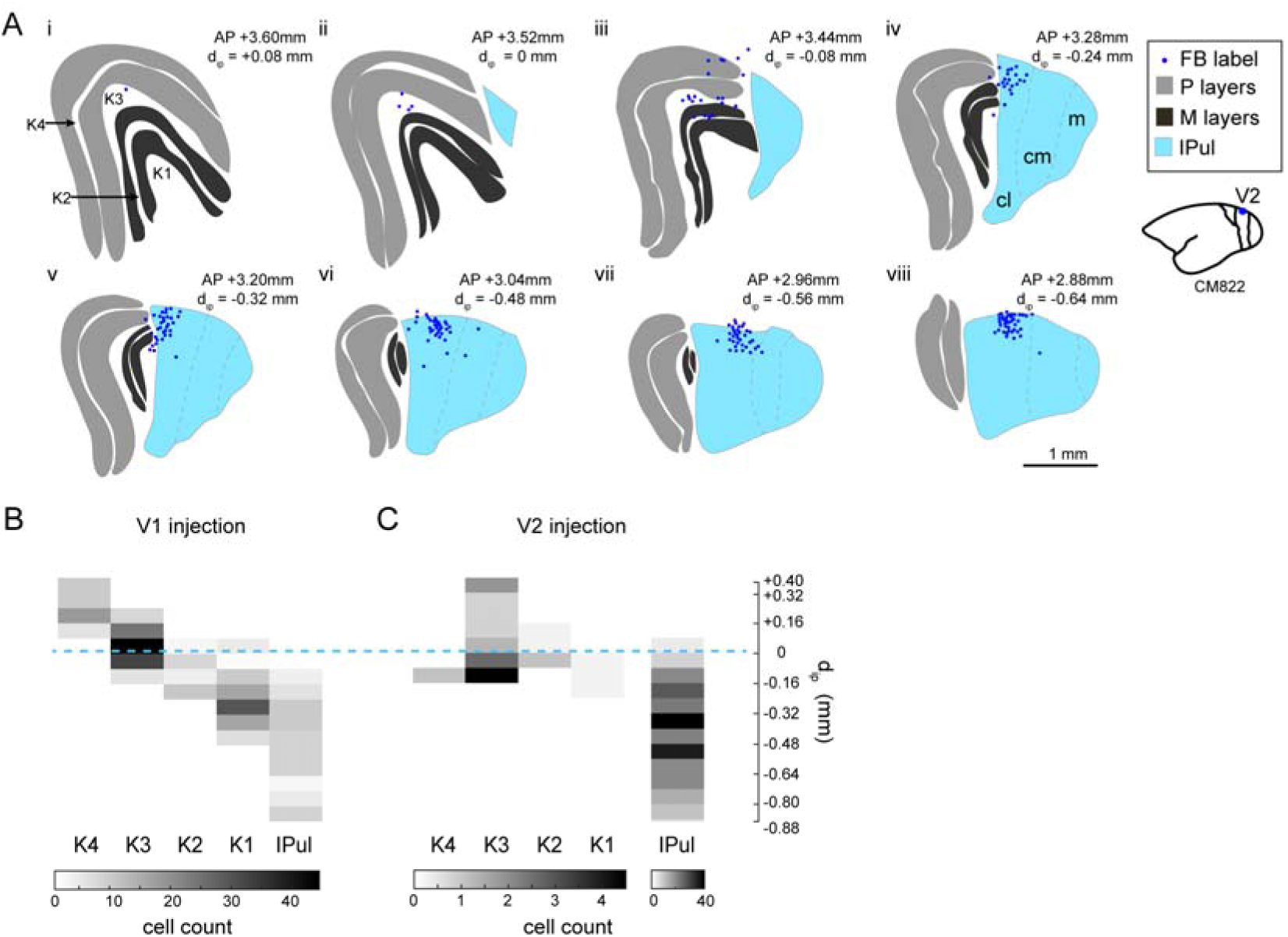
**A**. A series of annotated coronal sections showing LGN and IPul from rostral (i) to caudal (viii) direction for the case of M822 with FB injection in V2 (inset). FB-labeled cells (blue dots) were overlaid with the annotated brain regions. Subdivisions of IPul are indicated in (iv)-(vii). cl: caudolateral IPul; cm: caudomedial IPul; m: medial IPul. Other abbreviations are the same as Fig. 1. **B**. For all cases of retro-grade tracer injection in V1, the most rostral section with IPul appearance was set as the baseline section dip=0 (blue dashed line). A stack of coronal sections from dip = +0.4 mm anteriorly to dip = −0.88 mm posteriorly, were aligned at dip=0 for all cases. Each row represents one section, with 80 μm interval. The numbers of cells detected in K layers and IPul within each coronal section were represented with the darkness of each grid, where more cells correspond to darker grid. **C**. Similar to B, for all cases of retrograde tracer injection in V2, stacks of coronal sections were aligned by the most rostral section showing IPul (dip=0, blue dashed line). Number of cells in K layers and IPul within each section is represented in grayscale. Note that for V2 injection, the column of IPul is separated from the other regions due to the higher labeled cell density in IPul. Color scales are shown on the bottom.

All data analyses algorithms were written in Matlab (R2017b, Mathworks, USA) and Python 2.7.13. Image processing, cell detection, and fluorescent-to-Nissl registration were performed on GNU Linux operating on Intel(R) Xeon(R) 24-core CPU workstation. Other data analyses were performed on a Macintosh computer (Apple, Inc.).

## Results

### Calbindin-positive cells continued seamlessly from K layers of LGN to IPul

Calbindin staining revealed brightly labelled cells in both the LGN and IPul. In LGN, cells positive for calbindin were restricted to the K layers. The majority of calbindin positive cells were located in K1, K2 and K3. No clear cytoarchitectonic boundaries could be identified between LGN and IPul. The extension of calbindin-positive cells through the LGN K layers and IPul is demonstrated in Figure 1. Double labelling for Nissl substance (green) and calbindin (magenta) revealed the calbindin negative principal LGN laminae clearly (Fig.ure 1A). The calbindin positive cells in K layers and IPul formed a homogenous group.

### K layers and IPul form reciprocal connections with striate and extrastriate cortices

To visualize cortical connectivity in LGN K layers and IPul, anterograde and retrograde tracers were injected in the striate cortex (V1) and extrastriate cortices (V2 and DM). By projecting anterograde tracer-labelled axon terminal areas and retrograde-tracer labelled cell bodies from the fluorescent sections (Fig. 2B,E,H) to the brain region segmentation based on Nissl sections (Fig. 2A,D,G), we localized connections between LGN or IPul and visual cortex (Fig. 2C,F,I). The existence of each individual connection was reported as a percentage averaged across all cases (Fig. 3A, B), where 100% meant the existence of connection shown in all available cases. From anterograde tracing, V1 sends afferents to all layers of LGN and IPul. In 1 out of 3 cases, sparse labelling of terminals was found in K1 and IPul from V2 injection, but not any other LGN areas. In the rest 2 cases, V2 does not project to either LGN or IPul. DM projects to only K1 and IPul but not to other layers of LGN (Fig. 3A). From retrograde tracing, all layers of LGN and IPul send projections to V1. Retrograde tracer labelled cells from V2 and DM injections were found in IPul with high consistently and in LGN layers with different probabilities (Fig. 3B). Connection strengths from LGN layers and IPul to visual cortex were quantified with retrograde tracer-labelled cell counts, corrected by a profile-to-cell factor calculated based on detected profile area distribution (Fig. 3C,D). We observed strong connection from all layers of LGN to V1 (706±450 cells from P layers, 194±66 cells from M layers, and 589±502 cells from K layers). Projections towards V2 are rather inconsistent and sparse in P layers (50%, 1±1 cells) and M layers (50%, 3±3 cells), but much more consistent and denser from K layers (100%, 39±50 cells). From the only case of DM injection, we observed sparse labelling in all layers of LGN (9 cells in P layers, 2 cells in M layers, and 4 cells in K layers). Finally, IPul sends consistently strong afferents to V1 (174±94 cells), V2 (580±570 cells), and DM (295 cells).

### Retrograde labelled cells continued from K layers to lateral part of IPul

For both V1 and V2 injections, retrogradely labelled cells were detected more medially from anterior to posterior sections, corresponding to K4, K3, K2, K1 sequentially and eventually the lateral part of IPul. One example of this transition from V2 injection of FB is shown in Fig. 4A. In more anterior sections, sparse FB labelled cells were detected in K layers (Fig. 4A,a). With the sections more towards the brain posterior, FB labelled cells were detected more towards the medial part of the brain and the cell density increased (Fig. 4A, b-c). Eventually FB labelled cells were detected in both K1 and caudolateral IPul with higher cell density (Fig. 4A, d-e). Going further posterior, dense labelling of neurons were detected only in the caudolateral and caudomedial parts of IPul, but not LGN (Fig. 4A, f-h). This trend was observed in both V1 and V2 injections of retrograde tracers, and summarized for all cases in Fig. 4B. The sections were matched for all cases by aligning the most rostral section with IPul (blue dash line). We restricted the anterior-posterior range within +0.4 mm to −0.88 mm relative to this baseline section and calculated the average cell numbers per section within either IPul or K layers. The number of cells detected in each K layer and IPul were represented as grayscale for each section. For V1 injection, retrogradely labelled cells showed a clear transition from K4 caudomedially towards IPul (Fig. 4B). For V2 injection, K4 labelling was absent or sparse. From K3 to K1, retrogradely labelled cells were detected in more posterior sections. IPul labelling appeared in more caudal sections and was particularly dense compared to K layer labelling in the cases of V2 injections, plotted in a separate panel (Fig. 4C).

## Discussion

Similarities between LGN K layers and IPul in primates have long been noted (Xu *et al.*, 2001), yet few studies considered these two regions together (e.g. Warner *et al.*, 2012; Kwan *et al.*, 2018) since they were traditionally classified as parts of “first-order” and “higher-order” thalamic nuclei, respectively (Saalmann, 2014). In this study, we found that in marmosets, the calbindin positive cells in IPul and LGN formed a cytoarchitechtonically continuous population of cells, consistent with the previous observation in macaque (Jones & Hendry, 1989). We also found that cortical connections, either afferent or efferent, with K LGN layers and IPul always occur simultaneously, compared to the P and M LGN layers which lack projections from extrastriate cortices. Finally, the thalamic projection to V1 and V2 had a lateral to medial organization that crossed the borders between the LGN K layers and the lateral part of IPul. These results suggest that the K layers of LGN and IPul are anatomically continuous, and may underpin a single functional population of cells in this region of the thalamus, as discussed below.

In primates, calbindin labelling in the thalamus is associated with a role in cortical modulation and synchrony (Jones, 2007). Indeed, both the K LGN layers and vision-related pulvinar have been implicated in the modulation and synchronisation of cortical activity (Saalmann *et al.*, 2012; Klein *et al.*, 2016; Zhou *et al.*, 2016; Pietersen *et al.*, 2017). Similarities in function between the calbindin labelled K LGN cells and IPul cells in primate thalamus, as well as the observed continuity in calbindin labelling between the two structures are evidence that they make up a single functional population of thalamic neurons.

The continuity between K LGN cells and IPul also extends to the cortical connections of these thalamic structures. The most compelling evidence for this comes from the matched extrastriate inputs and outputs of the LGN K layers and IPul. Previous studies have reported V2 connections with K LGN cells and IPul independently (Hendry & Reid, 2000; Kaas & Lyon, 2007). In the current study we have shown that in all cases of V2 or DM anterograde tracer injection and V2 retrograde tracer injection, labelling was either seen in both LGN, especially K1, and IPul or not at all (Fig. 3A).

Subcortical inputs to the K LGN cells and IPul also provide further evidence that they form a single functional group of cells. Inputs from superior colliculus also showed “spill-over” of calbindin-positive cells from IPul to LGN (Jones, 2001). Separate studies on K cells and IPul revealed similar inputs from retinal ganglion cells: Kwan *et al.* (2018) demonstrated that retinal ganglion cells projecting to a sub-division of IPul in the marmoset had widefield morphologies including broad thorny, recursive bistratified, narrow thorny and large bistratified cells. Similar morphologies of retinal ganglion cells have been observed to project exclusively to the K layers of marmoset LGN (Szmajda *et al.*, 2008; Percival *et al.*, 2014; Masri *et al.*, 2017). Two visual pathways from retina to MT via K cells or IPul are alternatives from retino-geniculate-cortical pathway. They contribute to the early maturation of MT (Warner *et al.*, 2012), hence support the visual preservation after V1 lesion in early life, or “blindsight” (Warner *et al.*, 2015).

There are a few drawbacks of the current study. First, layer specificity of thalamocortical connections have been studied extensively (e.g. Jones 2009). However, in our tracing experiments, we filled the entire cortical column by releasing the tracer at different depths. As a result, no conclusion can be drawn from the current results regarding cortical layer specificity. Second, only 1 retrograde tracer injection was placed in DM, which limited our conclusion regarding the connection from DM to LGN and pulvinar. Previous studies have shown reciprocal connections between DM and IPul, but not LGN (Beck & Kaas, 1998). Here within the only case of DM injection, we observed very sparse labelling in LGN layers, including K layers. Further experiments are needed to confirm the existence of this connection.

## Acknowledgements

We thank Dr. Jaikishan Jayakumar for useful discussions on marmoset brain anatomy and fluorescent tracer detections.

## Conflict of interest

none

## Notes

Funding Information: National Health & Medical Research Council (NHMRC) Project grant (number 1081441 to PRM, UG); the Australian Research Council Centre of Excellence for Integrative Brain Function (CE140100007 to UG and PRM); Brain Mapping of Integrated Neurotechnologies for Disease Studies (Brain/MINDS) from the Japan Agency for Medical Research and Development, AMED (JP17dm0207001); National Institutes of Health (NIH) Grant 5R01EB022899.

